# iCNG99: a validated genome-scale metabolic model of *Cryptococcus neoformans strain* H99

**DOI:** 10.64898/2026.04.12.718001

**Authors:** Chun Feng, Pengjie Hu, Yin Zhu, Weixin Ke, Xindi Gao, Chen Ding, Bing Zhai, Linqi Wang, Ziwei Dai

## Abstract

*Cryptococcus neoformans* is a ubiquitous environmental fungus that can also cause life-threatening infections in immunocompromised individuals. As a competent pathogen, *Cryptococcus* needs to reprogram its metabolism to adapt to the drastic differences between environmental niches and host niches. A well-curated genome-scale metabolic model (GEM) is a powerful tool to facilitate the investigation of the metabolic resilience of an organism. Here we reconstructed and validated iCNG99, a GEM for *C. neoformans* reference strain H99, and evaluated its predictive performance across 43 growth conditions and gene essentiality benchmarks. The model achieved high confidence essential gene prediction (precision = 0.77) and recapitulated pathways targeted by clinically available antifungals. Integration with transcriptomic and metabolomic data enabled iCNG99 to capture condition-specific metabolic adaptations and to identify candidate vulnerabilities in drug tolerance, revealing metabolic adaptations associated with survival within host conditions and drug susceptibility. Together, iCNG99 provides a systems-level computational platform for investigating *C. neoformans* metabolism and for prioritizing antifungal vulnerabilities.

## Introduction

*Cryptococcus neoformans* is a globally distributed opportunistic fungal pathogen and a major cause of fungal meningitis, particularly in immunocompromised individuals such as those with advanced HIV/AIDS[1]. It is estimated to cause over 118,000 deaths annually, with the highest burden in sub-Saharan Africa, where limited access to timely diagnosis and effective antifungal therapy contributes to high mortality[2]. Infection is initiated through inhalation of environmental propagules, which establish in the lungs and can disseminate to the central nervous system, leading to life-threatening meningoencephalitis[3]. The virulence of *Cryptococcus* is supported by multiple traits, including a polysaccharide capsule that hinders host immune recognition, melanin production that protects against oxidative and thermal stress, and thermotolerance that enables survival in warm-blooded hosts[1, 7–9]. These features, combined with remarkable metabolic adaptability, enable its persistence in diverse and often hostile environments. Clinical treatment of cryptococcosis primarily relies on amphotericin B and a few azoles [10]. Changes in metabolic activity represent a key mechanism underlying microbial phenotypes and are closely linked to both virulence and drug resistance in *C. neoformans*.

In *C. neoformans*, metabolic research to date has largely focused on specific pathways, such as capsule production, energy metabolism and lipid metabolism, often in relation to virulence or stress adaptation[11–13]. While these studies have provided valuable biochemical insights, they remain fragmented and limited in scope, without an integrated framework to connect molecular changes across the whole metabolic network. Genome-scale metabolic models (GEMs) provide a systems-level framework for integrating genomic, biochemical, and physiological knowledge into comprehensive organism-wide metabolic networks[14]. When coupled with multi-omics datasets, they can elucidate how environmental perturbations, genetic modifications, or pharmacological interventions reshape metabolic states[15]. For example, in the research on other human fungal pathogens, such as *Candida albicans* and *Aspergillus fumigatus*, expertly curated GEMs that have facilitated systematic metabolic analyses and mechanistic links to infection biology[18–21]. For *Cryptococcus neoformans*, however, the absence of an equally well-curated GEM continues to hinder comprehensive investigation of its metabolism.

Here, we present iCNG99, a curated GEM for *C. neoformans* var. *grubii* H99 that incorporates key pathogen-specific pathways, including capsule biosynthesis and melanin production. The reconstruction combined semi-automatic computational pipeline with extensive manual curation to address challenges unique to non-model fungi. The reconstructed model adheres to current community standards and achieves a high MEMOTE score, supporting its structural completeness. We validated its predictive performance against growth phenotypes and gene essentiality data and further demonstrated its utility in pathogenicity-relevant conditions. Specifically, we applied iCNG99 to compare *in vivo* and *in vitro* metabolic states associated with fungal survival within host and to identify candidate antifungal targets in drug-tolerant states. Taken together, iCNG99 offers a computational platform for investigating the metabolic basis of *C. neoformans* pathogenicity and guiding antifungal target discovery.

## Methods

### Reconstruction of draft model

An initial draft genome-scale metabolic model (GEM) of *Cryptococcus neoformans* was constructed by integrating three KEGG-based reconstructions. Two drafts were generated using the RAVEN 2.0 toolbox[22]: one based on homology search with protein annotations from NCBI[23] RefSeq (GCF_000149245.1), and the other using species-specific KEGG[24] annotations (organism ID: cng). The third draft was reconstructed using Merlin[25], which combines homology-based annotation with the SamPler algorithm, using genome data from NCBI (taxonomy ID: 235443). Gene identifiers were standardized to RefSeq Protein Accessions using bioDBnet[26]. When gene-protein-reaction associations differed among drafts, they were merged using logical “or” to retain all gene assignments.

### Definition and calculation of biomass composition

Two separate biomass reactions were defined to represent the cell with and without capsule production, referred to as capsule-containing biomass and capsule-free biomass (Dataset S1), respectively. The biomass composition includes macromolecular fractions of DNA, RNA, protein, lipid, carbohydrate, and cofactors. These fractions were determined from experimental data and literature sources, with the calculation method adapted from the protocol by Thiele[27]. DNA, RNA, and protein precursors were estimated from genomic, transcriptomic, and proteomic sequences retrieved from NCBI RefSeq (GCF_000149245.1), and the values were weighted by molecular mass. The lipid composition, comprising free fatty acids, sterols, phospholipids, and sphingolipids, was derived from published data[28–30], and average molecular weights were calculated for fatty acids and sterols. Because the relative proportions of individual carbohydrate components in *C. neoformans* could not be determined from the literature, the carbohydrate composition was inferred from published data on the presence of these components in *C. neoformans* [31, 32]and by referencing the related model iSM996[33] from a species with a similar carbohydrate profile. For cofactors, which account for only a small fraction of the biomass, their possible presence in *C. neoformans* was first confirmed from the literature, and the specific composition was then directly adopted from the model iOD907[34]. The capsule which, not essential for growth but relevant for virulence, was included in capsule-containing biomass by assuming its mass accounts for 10% of the total carbohydrate content. The relative proportions of capsule monosaccharides were derived from analyses of the basic structural units of the two major polysaccharide components, glucuronoxylomannan (GXM) and galactoxylomannan (GalXM), as reported in the literature[35, 36]. In GXM, the molar ratio of xylose: mannose: glucuronic acid residues is 2:3:1, whereas in GalXM, the molar ratio of galactose: mannose: xylose: glucuronic acid residues is 6:4:2:1. These molar ratios were then used to calculate the corresponding mass fractions following the approach described above.

Growth-associated maintenance energy (GAM) was implemented through ATP hydrolysis reactions in the biomass formulation, using values referenced from the yeast model iMM904[37]. Non-growth-associated maintenance energy (NGAM) was introduced as a separate ATP-consuming reaction with a lower bound of 5 mmol gDW⁻¹, based on reported estimates in models of pathogenic yeast[19].

### Removal of redundancies and stoichiometric balancing

Redundant metabolites were removed by unifying KEGG entries with different identifiers for the same compound, retaining those with “C” prefixes. Overall multi-step reactions were deleted when all component steps were present. Ionic forms of metabolites at pH 7.2 were predicted using ChemAxon’s cxcalc (v19.13.0), and atomic composition and charge were assigned with RDKit[38]. A complete list of removed redundant metabolites is available in Dataset S2.

For reactions in which all participating metabolites had defined chemical structures, elemental and charge balancing was verified by comparing the total number of atoms and net charge on the reactant and product sides. When imbalances were detected, water and protons were used for adjustment where applicable. Reactions that could not be balanced in this way were resolved by consulting the MetaCyc database[39] and the PubChem database[40].

### Correction of the reversibility of the model

Reaction directionality was determined in two stages. In the first stage, reversibility and direction were assessed and, where necessary, corrected based on comparative analysis of pathway information from the KEGG Pathway and MetaCyc databases. For reactions present in both databases with consistent annotations, the shared directionality was adopted and labeled “MetaCyc; KEGG” in the model. When annotations differed, the MetaCyc assignment was used and labeled “MetaCyc-KEGG”. Reactions present in only one database were labeled “MetaCyc” or “KEGG”, respectively. If no directionality information was available in either database, relevant literature and other curated models were consulted, with annotations recorded as “PubMed” or “Other model”. In cases where no evidence was found, reactions were assumed to be reversible and labeled “None”.

After manual curation, standard Gibbs free energy changes (Δ G^O^) and thermodynamically feasible upper and lower bounds of reaction Gibbs free energy were computed using the dGbyG algorithm and thermodynamic flux balance analysis (TFBA) [41]. This analysis was used to evaluate whether reactions annotated as reversible could be thermodynamically constrained to irreversibility, and to verify the assigned directionality of reactions annotated as irreversible. Only reactions whose entire feasible Gibbs free energy range was ≤ −10 kJ mol⁻¹ or ≥ +10 kJ mol⁻¹ were considered for constraint adjustment.

### Gap-filling of the model

Exchange reactions were added to simulate nutrient uptake and metabolite secretion. A minimal YNB medium was defined using D-glucose as the sole carbon source and ammonium sulfate as the sole nitrogen source, with bounds set to −1000 and 1000. Additional exchange reactions were added for other carbon and nitrogen sources tested experimentally (Dataset S3), with uptake disabled (lower bound = 0) when not in use. Growth simulations were performed using COBRApy[42] by sequentially setting each biomass component as the objective function. For conditions known to induce capsule production, the capsule-containing biomass reaction was used. In conditions where capsule formation was absent or uncertain, the capsule-free biomass formulation, was applied. If no flux was observed, indicating a gap in the biosynthetic pathway, reactions were added based on KEGG, MetaCyc, literature, and other published models to restore connectivity, while minimizing the number of added reactions. Growth under alternative carbon and nitrogen sources was also tested, and gap-filling was performed in cases where model predictions were inconsistent with experimental results. In addition, melanin biosynthesis was simulated under conditions with phenol supplied in the medium, and the relevant reactions[9] were incorporated into the model.

### Assign compartments and transport reactions

Protein subcellular localization was predicted using an integrative approach combining results from three multi-label predictors: Wolf Psort[43], DeepLoc 2.0[44], and YLoc[45]. For Wolf Psort, compartments with scores ≥ 80% of the highest were retained. Prediction scores were set to 1, except for low-confidence YLoc results, which were weighted as 0.5. Final localization was assigned by summing scores across tools; compartments with total scores ≥ 2 were selected, or ≥ 1.5 if no higher score was available. Proteins without confident predictions were assigned to the cytoplasm. Only eight shared compartments were considered; others, including the plasma membrane, were reassigned to the cytoplasm. Reactions were localized based on associated enzymes, or to the cytoplasm if no gene association was present. The final model included seven compartments.

Based on the predicted compartment information, transport reactions were integrated from three sources: predictions from the TranSyt toolbox[46], the yeast consensus model version 8.7[47], and manually defined reactions. All transport reactions predicted by TranSyt were directly incorporated into the model. These reactions were generated using a TC family threshold of 0.7, with ontology filters disabled, and reassigned to appropriate compartments based on predicted protein localization and metabolite distribution. Reactions from the yeast model were selected by matching metabolites and compartments, treating organelle membranes as part of the corresponding organelle. Manually defined reactions were added between the cytoplasm and other compartments for shared metabolites. For candidate transport reactions not derived from TranSyt, a mixed-integer linear programming (MILP) model was formulated to select a minimal reaction set. Each candidate reaction was assigned a binary decision variable and weighted according to its source, with yeast model reactions assigned a weight of 1 and manually defined reactions assigned a weight of 10. The MILP was solved under 43 carbon and nitrogen source conditions, with the constraint that the biomass production rate must reach at least 10% of that obtained using the complete transport reaction set. The union of reactions selected across all conditions was then incorporated into the final model.

### Database cross-referencing for genes, metabolites, and reactions

Gene annotations were supplemented with identifiers from NCBI Gene, NCBI Protein, and UniProt[48], obtained via the KEGG database. For metabolites and reactions, MetaNetX[49] was used to add identifiers from PubChem, MetaNetX, BiGG[50], HMDB[51], ModelSEED[52], and Reactome[53]. Additionally, reactions were further annotated with BioCyc and Systems Biology Ontology (SBO) identifiers.

### Assessment of model structural quality

Model quality was first evaluated using MEMOTE[54], which provides a standardized assessment of metabolic model structure and annotation. Additional consistency checks were performed using COBRA[55], including a leak test to verify that no metabolite production occurs when all uptake reactions are closed. Simulations were also conducted to test whether protons could be produced from water or oxygen alone, and whether any biomass precursors or energy-containing metabolites could be generated when only ATP hydrolysis was allowed. These tests were used to identify and eliminate thermodynamically infeasible or mass-violating reactions.

### Gene essentiality simulation

The gene encoding the known target of an azole antifungal (ERG11), identified as CNAG_00040, was retrieved from UniProt. Single-gene deletion was simulated by removing the target gene and modifying the flux bounds of associated reactions according to the gene–protein–reaction (GPR) rules. For reactions regulated by the gene through an “AND” relationship or as the sole gene contributor, both the upper and lower bounds were set to zero. The model was then re-optimized under standard growth conditions to record biomass flux and evaluate the impact of gene deletion.

After validating the single drug target, large-scale single-gene knockout simulations were performed under YPAD medium conditions. Single gene knockouts were considered essential if the simulated biomass flux dropped to zero. Flux balance analysis was used to compute the optimal biomass flux for each knockout mutant. Model-predicted essential and non-essential genes were compared against genome-wide transposon insertion screening data[56] to construct a confusion matrix. Accuracy, precision, recall, F1 score, and Matthews correlation coefficient (MCC) were calculated to evaluate prediction performance.

For model false negative (FN) genes, which are those predicted as non-essential but experimentally found to be essential, functional annotation and enrichment analyses were conducted using KEGG pathway enrichment and GO biological process categories. Significance was determined by Benjamini-Hochberg corrected q-values with a threshold of < 0.05. Protein sequences of FN genes were aligned against the MoonProt database using BLAST with an E-value threshold of < 1 × 10^-5^ to identify potential moonlighting proteins.

### Culturing conditions

*C. neoformans* strain H99 was cultured in YPD medium (10 g/L yeast extract, 20 g/L peptone, 10 g/L glucose) for 16 hours. Cells were harvested by centrifugation and washed twice with sterile water. The washed cells were inoculated into YNB medium (1.7 g/L yeast nitrogen base, 5 g/L ammonium sulfate) supplemented with 20 g/L of various carbon sources, including glycerol, D-trehalose, succinic acid, erythritol, and D-melibiose and so on (Dataset S3). Cultures were incubated in a multi-mode microplate reader (CLARIOstar Plus/FLUOstar Omega/ACU) at 30°C with shaking at 600 rpm for 48 h. After that, capsule formation was visualized by staining the cells with India ink and observing them under a microscope.

### RNA-seq

Wild-type *C. neoformans H99 strain* at stationary phase were generated by transferring colonies grown on YPD agar for 24 h into 3 mL liquid YPD and incubating at 30 °C on a slanted shaker set at a 45° angle for 12 h. Cultures were then re inoculated into 4.0 mL fresh liquid YPD at an initial density of 1 × 10^7^ cells/mL and incubated under the same conditions for 72 h to obtain stationary phase cells. For infection, cells were washed twice with phosphate buffered saline by centrifugation at 4000 rpm for 2 min, resuspended in 1 mL sterile phosphate buffered saline, and counted using a hemocytometer. The inoculum was adjusted to 2 × 10^8^ cells/mL, and mice were anesthetized with isoflurane and intranasally inoculated with 50 μL of the suspension, corresponding to 1.0 × 10^7^ cells per mouse. Animals were maintained on a warming pad until recovery and then returned to their cages. At 7 days post infection, lungs were harvested, snap frozen in liquid nitrogen, and processed for total RNA extraction. RNA sequencing was performed by Novogene (Beijing, China).

### RNA-seq data preprocessing

Raw RNA-seq data for *in vivo* and *in vitro* conditions were obtained from two independent sources, including publicly available datasets from the GEO database (GEO accession: GSE122785)[57] and experiments conducted in this study. In addition, transcriptomic data for drug-tolerant and non-drug-tolerant strains (GEO accession: GSE188965)[58], as well as thermotolerant and non-thermotolerant strains (GEO accession: GSE242109) were retrieved from the GEO database. The corresponding reference genome was retrieved from NCBI (accession number: GCF_000149245.1). Quality control of the sequencing data was performed using FastQC[59], followed by adapter trimming and removal of low-quality reads with Trim Galore[60]. Paired-end alignment was conducted using STAR v2.7.11b[61], and output files were converted with SAMtools[62]. Read counting was performed using featureCounts[63], focusing on reads mapped to exonic regions and excluding multi-mapped reads.

### Transcriptomic correlation analysis

Log_2_ fold changes (log_2_FC) in gene expression between *in vivo* and *in vitro* conditions were independently calculated using DESeq2 for two transcriptomic datasets. Genes shared between the two datasets, as well as metabolic genes included in the model, were identified, and their corresponding log_2_ fold change (log_2_FC) values were extracted. For both gene sets, Spearman’s rank correlation coefficients were calculated to assess the transcriptional correspondence between the two data sources.

### Condition-specific model construction

RNA-Seq raw read counts were normalized to transcripts per million (TPM) to obtain comparable gene expression levels across samples. Gene-level expression values were mapped to metabolic reactions according to gene-protein-reaction associations, using the arithmetic mean for “or” relationships and the geometric mean for “and” relationships. Reaction-level regulation was assessed by calculating log₂ fold changes between the studied conditions based on the mapped reaction expression values. Reactions were classified as upregulated or downregulated if they met the criteria of an adjusted p-value < 0.05 and an absolute log_2_ fold change exceeding a condition-specific threshold.

Flux variability analysis (FVA) was performed in COBRApy under the relevant nutrient constraints to determine the set of reactions active in each condition, with reactions considered active if the absolute value of their minimum or maximum feasible flux exceeded 1×10⁻□mmol·gDW⁻¹·h⁻¹. The intersection of active reactions and significantly upregulated or downregulated reactions was used as input to an improved REMI algorithm [64], which integrates transcriptomic constraints into the base metabolic model to generate a relative, condition-specific genome-scale metabolic model.

For the thermotolerance analysis, raw metabolomics data were obtained from the OMIX, China National Center for Bioinformation/Beijing Institute of Genomics, Chinese Academy of Sciences (https://ngdc.cncb.ac.cn/omix) with the accession numbers: OMIX010912. Metabolites were classified as upregulated or downregulated if they met the criterion of an adjusted p-value < 0.05 and an absolute log₂ fold change ≥ 0.585. Dead-end metabolites were excluded from the analysis.

### Flux distribution and enrichment analysis

For each condition-specific model, the REMI algorithm was used to find all alternative optimal flux distributions. Reactions with the same flux direction across all solutions and showing consistent upregulation or downregulation in more than 80% of the alternative solutions were selected, and the geometric mean of their log_2_ scaled flux changes was calculated. These reactions, excluding sink and exchange reactions, were then subjected to model-based pathway enrichment analysis to identify significantly perturbed metabolic modules, and flux correlation analysis comparing log_2_ scaled flux fold changes between the two datasets derived from independent sources.

### Essential gene prediction in AmB-tolerant state

Context-specific models for the AmB-tolerant and non-tolerant states were constructed using the GIMME algorithm in the COBRA toolbox based on their respective transcriptomic profiles. Single-gene knockout simulations were performed for each model to identify essential genes under the given condition. Genes were considered essential if their deletion resulted in a predicted growth rate of zero.

Essential genes specific to the drug-resistant state were compared against the human proteome using the BLASTP algorithm [64]. A gene was considered non-homologous to human proteins if no BLASTP hit met the significance criteria of sequence identity ≥ 70% and an E-value ≤ 1 × 10⁻□. Genes classified as non-homologous were selected as potential drug targets.

### Growth measurement of *C. neoformans* gene deletion strains

Wild-type strain and gene deletion strains were performed under the drug-tolerant state in glucose-containing medium at 30 °C with shaking. Cultures were set up in 96-well plates with a working volume of 1 mL per well. The initial inoculum was adjusted to an OD600 of 0.01. Growth was assessed by measuring the OD_600_ at 48 h post-inoculation and statistical significance was evaluated by Dunnett’s post-hoc test in R.

## Results

### Reconstruction of GEM for *C. neoformans* var. *grubii* H99

We developed and curated a genome-scale metabolic model of *Cryptococcus neoformans*, named iCNG99 (Fig. 1). To comprehensively capture the metabolic functions of *C. neoformans*, we integrated multiple draft reconstructions derived from different strategies into a more extensive draft network (Supplementary Fig. 1A). Compared to single-source drafts, this network showed broader gene and metabolite coverage but also introduced more blocked reactions and dead-end metabolites (Supplementary Fig. 1B, 1C), highlighting the need for systematic refinement to ensure functionality and biological relevance (Fig. 1A). During subsequent optimization, we employed an ensemble-based approach that combines multi-tool predictions for subcellular localization of reactions and integrated information from multiple databases to curate subcellular localization of reactions and transport reactions (Methods, Fig. 1B). By adopting a minimal set of transport reactions (Methods), we ensured a biologically coherent distribution of metabolites within the cell, while avoiding redundant or uncertain transport routes that could otherwise cause spurious leakage. We further defined two biomass reactions corresponding to capsule-containing and capsule-free states, respectively. The final formulation encompassed 103 precursors across DNA, RNA, proteins, lipids, carbohydrates, and cofactors (Fig. 1C; Dataset S1), representing a level of refinement that exceeds existing fungal reconstructions [16]. In particular, the lipid fraction comprises 55 lipid species, notably sphingolipids and phospholipids, which are crucial for maintaining membrane integrity and influencing pathogenicity.

**Figure 1:**
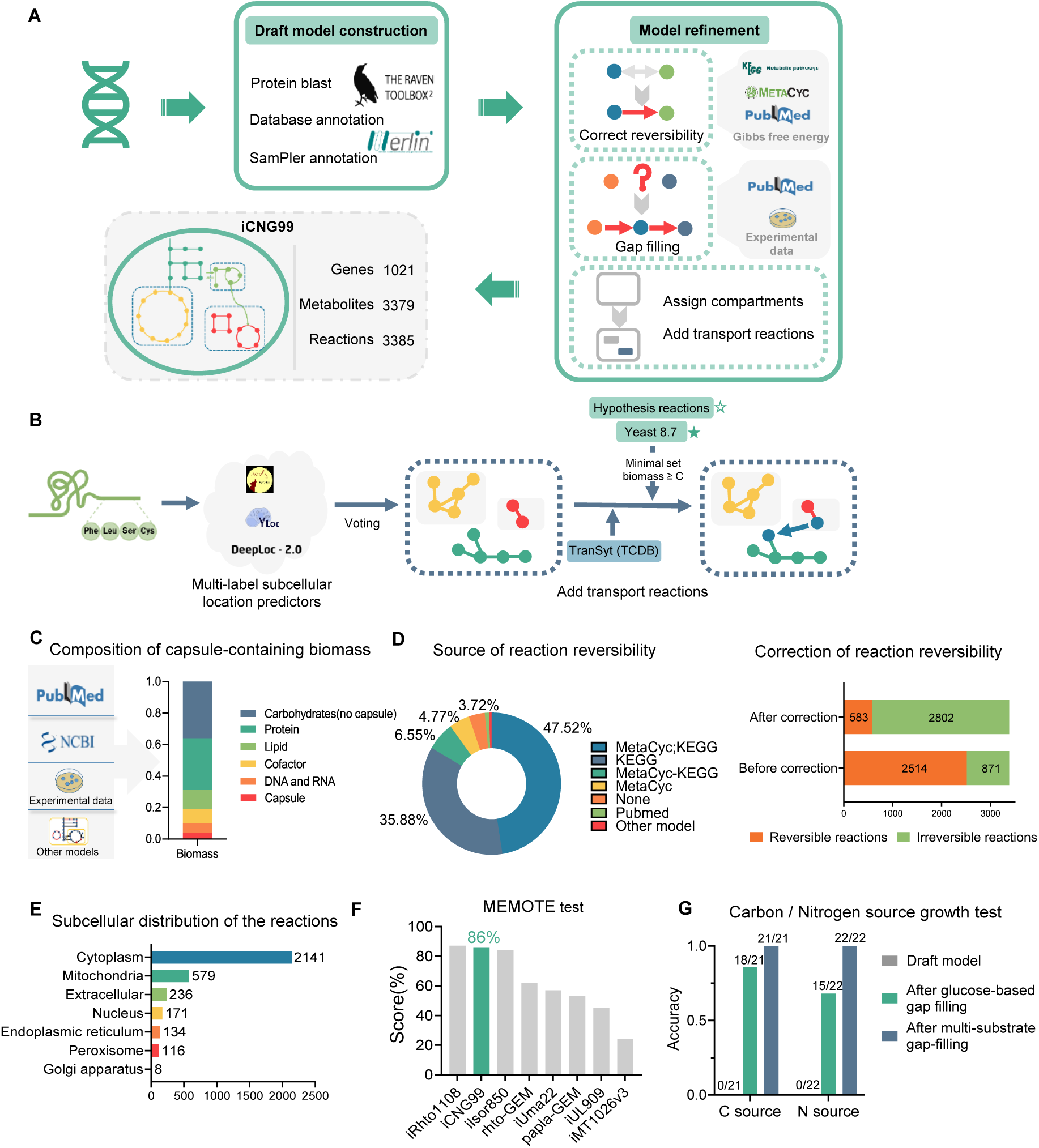
Construction and basic information of iCNG99. A. Workflow for construction of the genome-scale metabolic network model iCNG99 of *Cryptococcus neoformans*. B. Ensemble-based determination of subcellular location and transport reactions. C. Composition of the capsule-containing biomass reaction. D. Sources of information used in reaction directionality refinement and proportions of reversible and irreversible reactions before and after correction. “MetaCyc; KEGG” indicates reactions with consistent directionality in both databases, whereas “MetaCyc–KEGG” denotes discrepancies, with directionality assigned according to MetaCyc. E. Distribution of reactions across seven subcellular compartments. F. MEMOTE scores of iCNG99 and other recent fungal models. G. Accuracy of model-predicted growth across 43 experimentally tested carbon and nitrogen sources in the draft model, after glucose-based gap filling, and after multi-substrate gap filling.

Next, we combined information from KEGG, MetaCyc, published fungal models, and literature (Fig. 1D), followed by thermodynamic analysis of the network to constrain the reaction directionality (Methods). After correction, approximately two-thirds of reactions were assigned with confident directionality. In addition, we performed charge and atom balancing, identified and removed redundant metabolites and reactions, and incorporated cross-references from multiple databases into the model (Methods, Dataset S2) to enhance its biochemical consistency. The final reconstruction, named iCNG99, comprised 1021 genes, 3379 metabolites, and 3385 reactions distributed across seven compartments, including the cytoplasm, mitochondria, Golgi, vacuole, nucleus, peroxisome, and extracellular space (Fig. 1E).

### Validation of the iCNG99 model

To ensure that the reconstructed model was both structurally reliable and biologically predictive, we subjected iCNG99 to systematic validation. MEMOTE testing yielded a score substantially higher than most up-to-date fungal models, underscoring the improved annotation quality and network completeness (Fig. 1F). Further structural evaluation through leakage testing confirmed the absence of non-physiological energy cycles or metabolite loops, ensuring that the model does not harbor common topological flaws. Together, these results indicate that the reliability of iCNG99 is comparable to that of well-curated genome-scale reconstructions, providing a robust basis for subsequent functional validation.

We next assessed the predictive performance of iCNG99 against growth under different carbon and nitrogen sources. After iterative gap-filling guided by MetaCyc and KEGG pathways and cross-referencing with fungal models, iCNG99 successfully recapitulated growth across all 43 tested carbon/nitrogen conditions (Fig. 1G, Dataset S3) and non-growth under lignin and several D-amino acids, in agreement with wet-lab observations[65, 66]. This marked improvement highlights the capacity of iCNG99 to capture growth phenotype of *C. neoformans* with high specificity.

To evaluate the reliability of iCNG99 in predicting gene essentiality, we first selected the clinically validated azole target *ERG11* as a representative case for functional consistency validation (Fig. 2A). Under YPAD conditions, constraining the flux through *ERG11*-associated reactions to zero led to a complete loss of predicted growth (Fig. 2A), consistent with the known mechanism whereby inhibition of *ERG11* disrupts ergosterol biosynthesis and membrane integrity, causing growth arrest.

**Figure 2.**
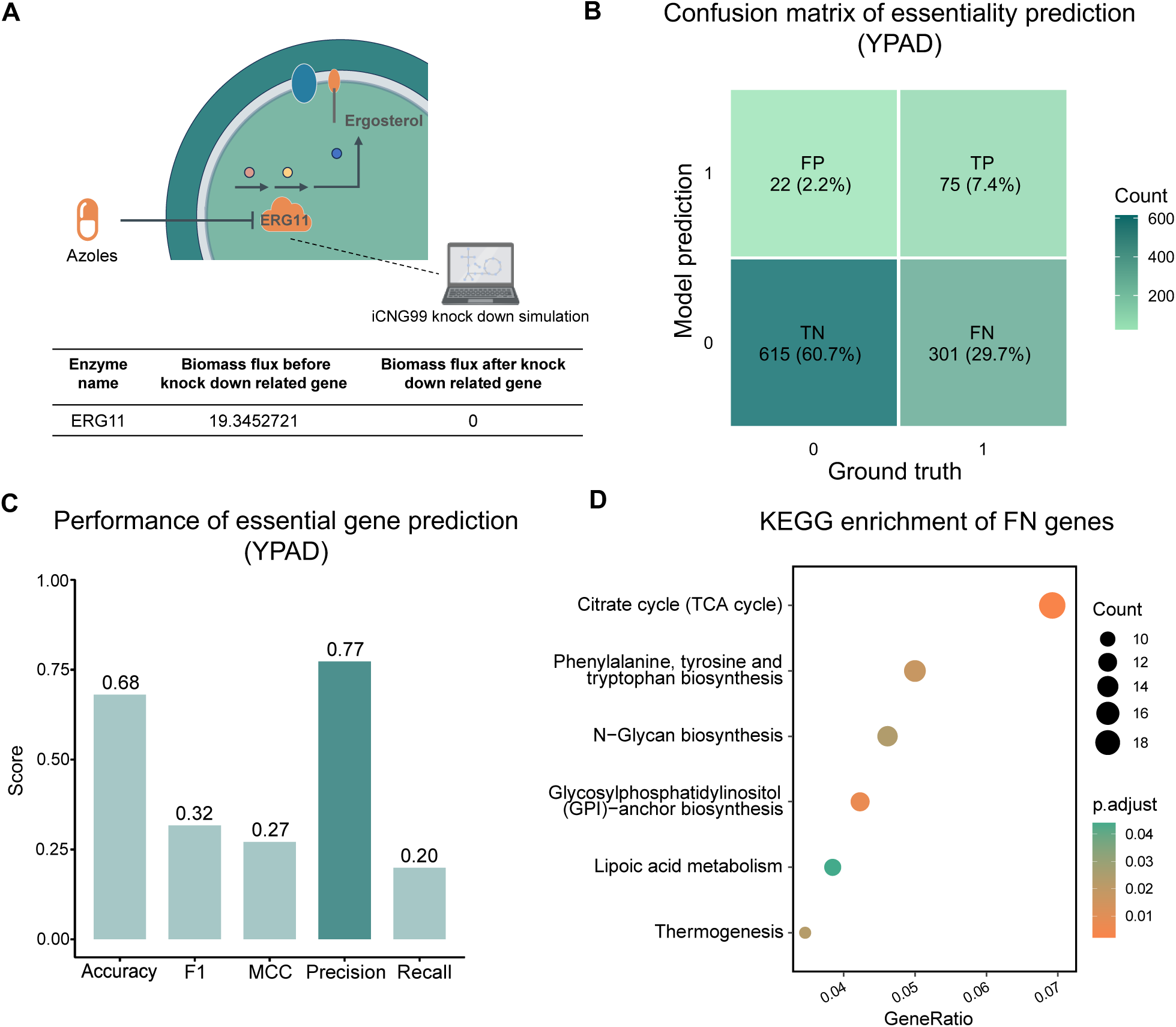
Validation of model-predicted essential genes. A. Model-predicted impact of ablating the known antifungal drug target ERG11 on the biomass flux. B. Confusion matrix comparing model-predicted gene essentiality with experimental results under YPAD culture conditions. 0: non-essential gene; 1: essential gene. C. Performance metrics of gene essentiality prediction by the iCNG99 model under YPAD conditions. D. KEGG pathway enrichment analysis of false-negative (FN) genes in the essentiality prediction.

We next systematically benchmarked iCNG99’s genome-scale essentiality predictions against a large-scale transposon deletion library (Fig. 2B). Under YPAD conditions, 1,013 genes were matched between the model and experimental dataset, yielding 75 true positives (TP), 615 true negatives (TN), 22 false positives (FP), and 301 false negatives (FN). The resulting confusion matrix (Fig. 2C) yielded an overall accuracy of 0.68, F1 score of 0.32, and Matthews correlation coefficient (MCC) of 0.27. For the essential class, the precision reached 0.77 with a recall of 0.20. Thus, 77% of genes predicted as essential by iCNG99 were supported by experimental evidence.

To investigate the causes of low recall, we examined the FN genes set through functional enrichment and multi-function analysis (Fig. 2D, Supplementary Fig 2, Dataset S4). KEGG and GO analyses showed significant enrichment in functions beyond core metabolism, involving protein modification, complex assembly, and organelle homeostasis, suggesting that causes for essentiality of these genes are not fully represented in current GEM formations (Fig. 2D, Supplementary Fig 2). BLAST searches against the MoonProt[67] database identified 73 proteins among the 301 FN genes sharing significant similarity with known moonlighting proteins (Dataset S4), suggesting that some FN genes may encode multi-functional proteins with both metabolic and non-metabolic roles. Moreover, since most GPR relations in iCNG99 use logical “OR” operators and rarely impose “AND” constraints for multi-subunit complexes or paralogs, these simplifications collectively contribute to a systematic underestimation of gene essentiality. Nevertheless, the high precision indicates that iCNG99 is particularly suitable for high confidence target identification.

### Integration of iCNG99 and transcriptomics reveals consistent *in vivo* metabolic reprogramming across independent datasets

GEMs are widely applied to integrate omics data for condition-specific metabolic modeling. However, transcriptomic datasets are often affected by noise and batch effects, which may compromise the stability and reliability of downstream metabolic predictions. To assess the reproducibility and interpretability of predictions generated from iCNG99, we obtained two independently derived RNA-seq datasets for both *in vivo* and *in vitro* conditions and evaluated whether consistent metabolic phenotype could be captured across conditions. We constructed condition-specific models using an optimized REMI workflow that directly integrates transcriptomics into iCNG99 (Fig. 3A, Methods). Transcriptomic datasets from both conditions were uniformly processed through the same pipeline for quality control and differential expression analysis (Methods). REMI-based consistency constraints were then applied as a core step, ensuring that only those reactions with flux directions concordant with transcript-level changes remained active. This strategy eliminated structural biases inherent to single-condition reconstructions and emphasized genuine metabolic shifts between environments.

**Figure 3.**
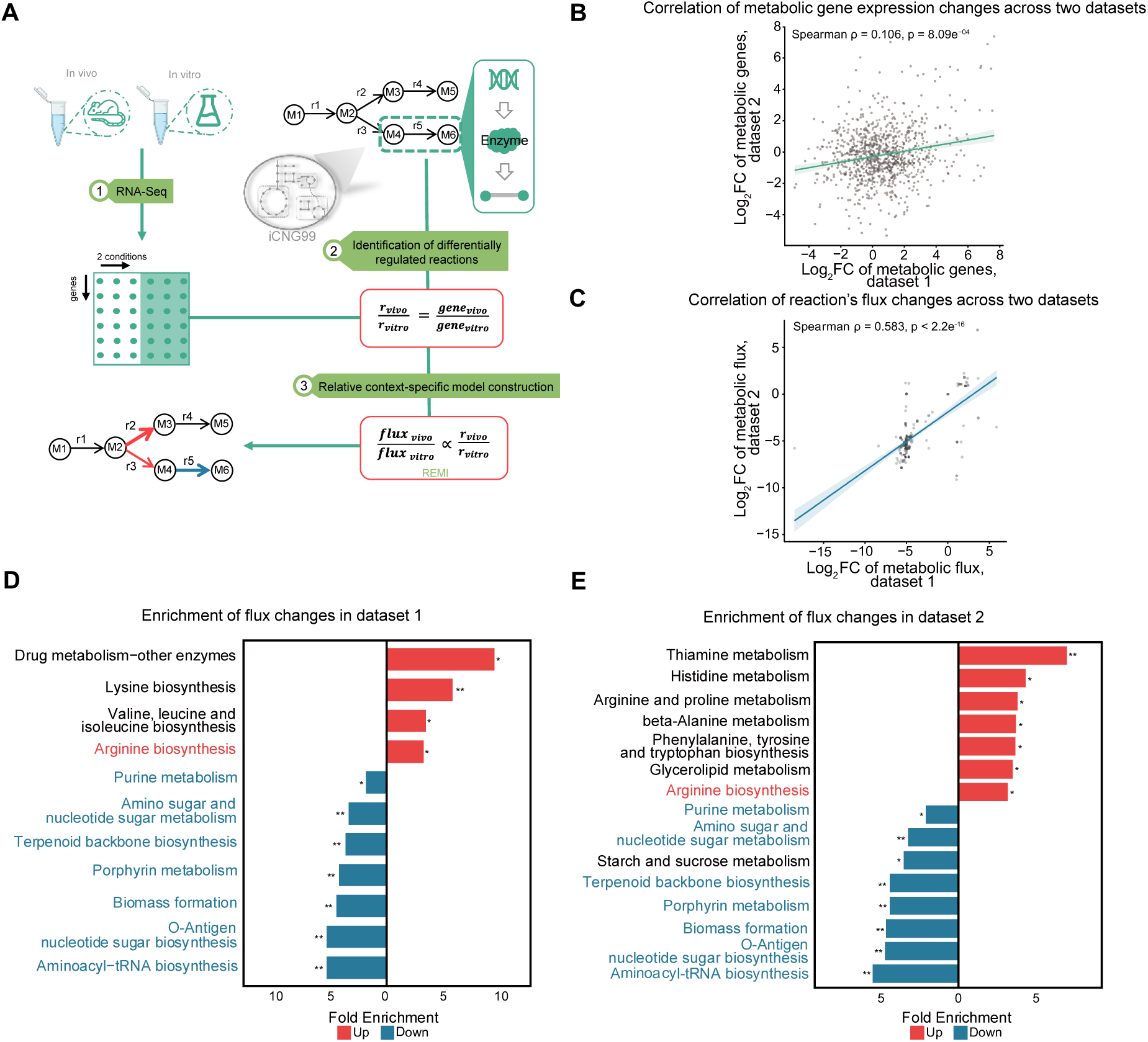
iCNG99 predicts metabolic changes reproducible between independent datasets. A. Workflow for constructing the condition-specific model by integrating omics data. B. Correlation of log_2_ fold changes in metabolic gene expression (*in vivo* versus *in vitro*) between two independent transcriptomic datasets. C. Correlation of log_2_ fold changes in model-predicted metabolic fluxes (*in vivo* versus *in vitro*) between two independent transcriptomic datasets. D. Pathways enriched in significantly altered metabolic fluxes between in vivo and in vitro conditions, dataset 1 (* p.adjust < 0.05, ** p.adjust < 0.01). E. Pathways enriched in significantly altered metabolic fluxes between *in vivo* and *in vitro* conditions, dataset 2 (* p.adjust < 0.05, ** p.adjust < 0.01). Colored pathway names indicate pathways significantly enriched in both datasets.

Although correlation in transcriptomic changes was weak between the two independent datasets (Spearman’s rank correlation coefficient = 0.106, Fig. 3B), integration of the transcriptomic datasets into context-specific metabolic models increased the correlation at flux level (Spearman’s rank correlation coefficient = 0.584, Fig. 3C, Supplementary Fig. 3). The concordance in metabolic changes between the two datasets was further supported by consistent directional changes across multiple metabolic modules (Fig. 3D, 3E). Amino acid-related pathways, particularly arginine biosynthesis, were consistently enriched among upregulated fluxes in both datasets. In contrast, pathways associated with biomass formation, aminoacyl-tRNA biosynthesis, purine metabolism, O-Antigen nucleotide sugar metabolism, terpenoid backbone biosynthesis, and porphyrin metabolism showed concordant downregulation across conditions.

Overall, these results indicate that, despite the presence of batch effects between transcriptomic datasets from independent sources, the integration into iCNG99 significantly enhanced the robustness and reproducibility of metabolic flux predictions. This concordance reveals a robust metabolic adaptation upon *in vitro* to *in vivo* transition, characterized by a shift from growth-supporting metabolic pathways toward cellular maintenance and adaptive regulation.

### Integrative modeling identifies membrane-centered metabolic remodeling under heat stress

In addition to the metabolic adaptation associated with the transition from *in vitro* to *in vivo* conditions, *C. neoformans* must also adapt to elevated temperature as a major challenge encountered in the host environment. To further evaluate the ability of the iCNG99 model to characterize metabolic reprogramming under this condition, we integrated transcriptomic and metabolomic data and computed reaction flux changes across different thermal conditions (Fig. 4). Heat stress is an important factor for *C. neoformans* in adapting to host temperature, particularly during pathogenicity, as changes in host body temperature significantly affect the growth and virulence of the fungus[8]. In response to elevated temperatures, *C. neoformans* must undergo metabolic reprogramming to adapt to the host’s high-temperature environment, ensuring its growth and survival. Transcriptomic enrichment analysis of the wild-type strain revealed upregulation of the MAPK signaling pathway and downregulation of ribosome-related genes (Fig. 4A), indicating that heat stress triggers canonical signaling activation coupled with translational repression. However, the integrated modeling uncovered additional metabolic-layer remodeling that was not evident from expression data alone (Fig. 4B). The predicted flux distribution showed a marked convergence toward sphingolipid metabolism, whereas valine, leucine and isoleucine degradation, porphyrin metabolism, glycerophospholipid metabolism, and inositol phosphate metabolism were downregulated. This reprogramming of metabolic fluxes reflects a shift from energy-consuming pathways toward membrane lipid biosynthesis, consistent with the membrane remodeling observed during thermal adaptation[68].

**Figure 4:**
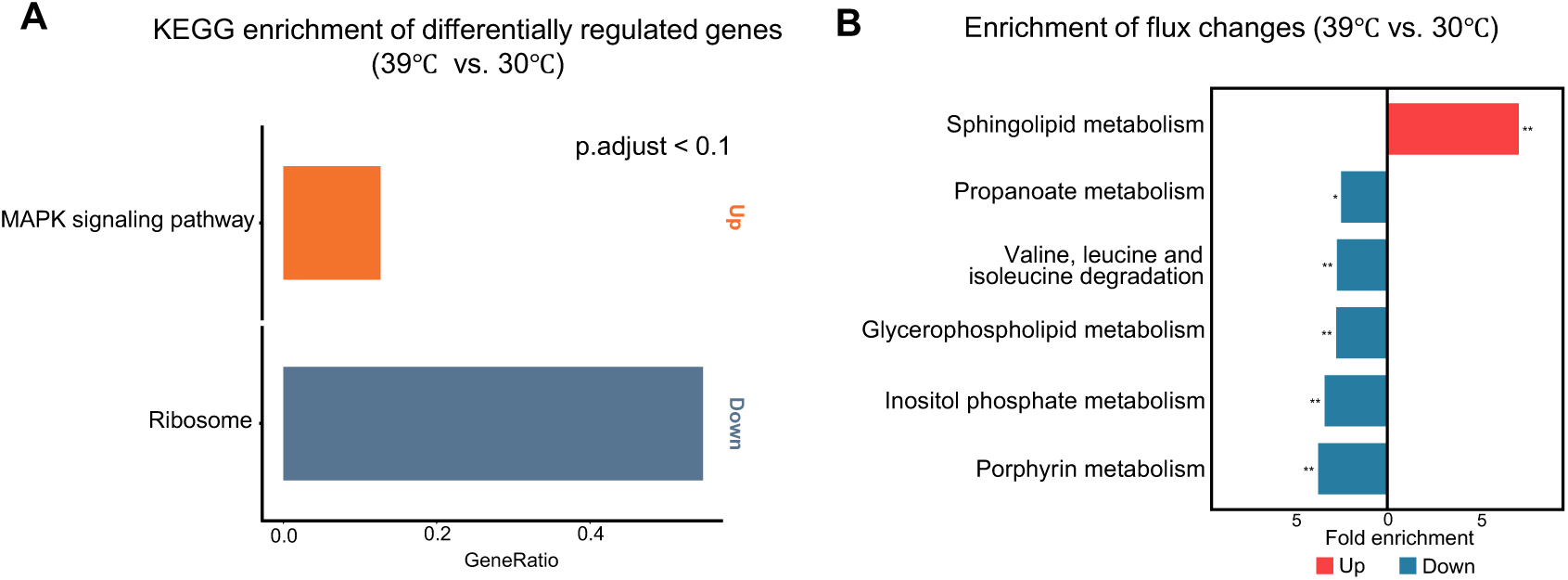
iCNG99 predicts metabolic shifts in response to heat. A. Pathways enriched in differentially expressed genes at 39℃ versus 30℃ conditions. B. Pathways enriched in metabolic fluxes altered at 39℃ versus 30℃ conditions predicted by the condition-specific GEMs (* p.adjust < 0.05, ** p.adjust < 0.01).

### Condition-specific essentiality analysis identifies potential metabolic vulnerabilities in amphotericin B tolerance

Another important application of GEMs is in network pharmacology, particularly in identifying potential drug targets. Drug tolerance is a key factor in the treatment failure of *C. neoformans*[69], and metabolic pathways are at the core of fungal function and physiological expression, directly influencing how the fungus responds to environmental changes and therapeutic pressures. By integrating the iCNG99 model with transcriptomic data in the state of tolerance to amphotericin B[58], we constructed condition-specific metabolic model for *C. neoformans* under drug tolerance and performed essentiality analysis (Fig. 5A). The drug-tolerant state model predicted 106 essential genes, forming the core metabolic backbone required for cellular viability (Fig. 5B, Dataset S5). Cross-species homology screening further identified 24 strain-specific essential genes that lacked human orthologs and were predominantly enriched in lipid metabolism, as well as amino acid, carbohydrate, and nucleotide metabolism pathways (Fig. 5C). Comparison with the condition-specific knockout mutant library[8, 70] revealed that 18 of these genes lacked corresponding deletion strains, suggesting their potential essentiality for fungal growth under the tolerant state (Fig. 5B). For the remaining 6 genes with available knockout mutants, growth was assessed by measuring OD_600_ at 48 h post-inoculation (Fig. 5D). 5 mutant strains exhibited significantly reduced growth compared with the wild type at 48 hours, while one gene showed a marginally significant decrease (p.adjust = 0.073). These experimental observations were highly consistent with our model predictions, demonstrating the accuracy and biological relevance of the model in predicting metabolic essentiality under the drug-tolerant state. Collectively, the 18 genes constitute a preliminarily validated set of strain-specific antifungal target candidates, with lipid-related functions emerging as dominant metabolic vulnerabilities that may inform the rational design of therapeutic strategies.

**Figure 5.**
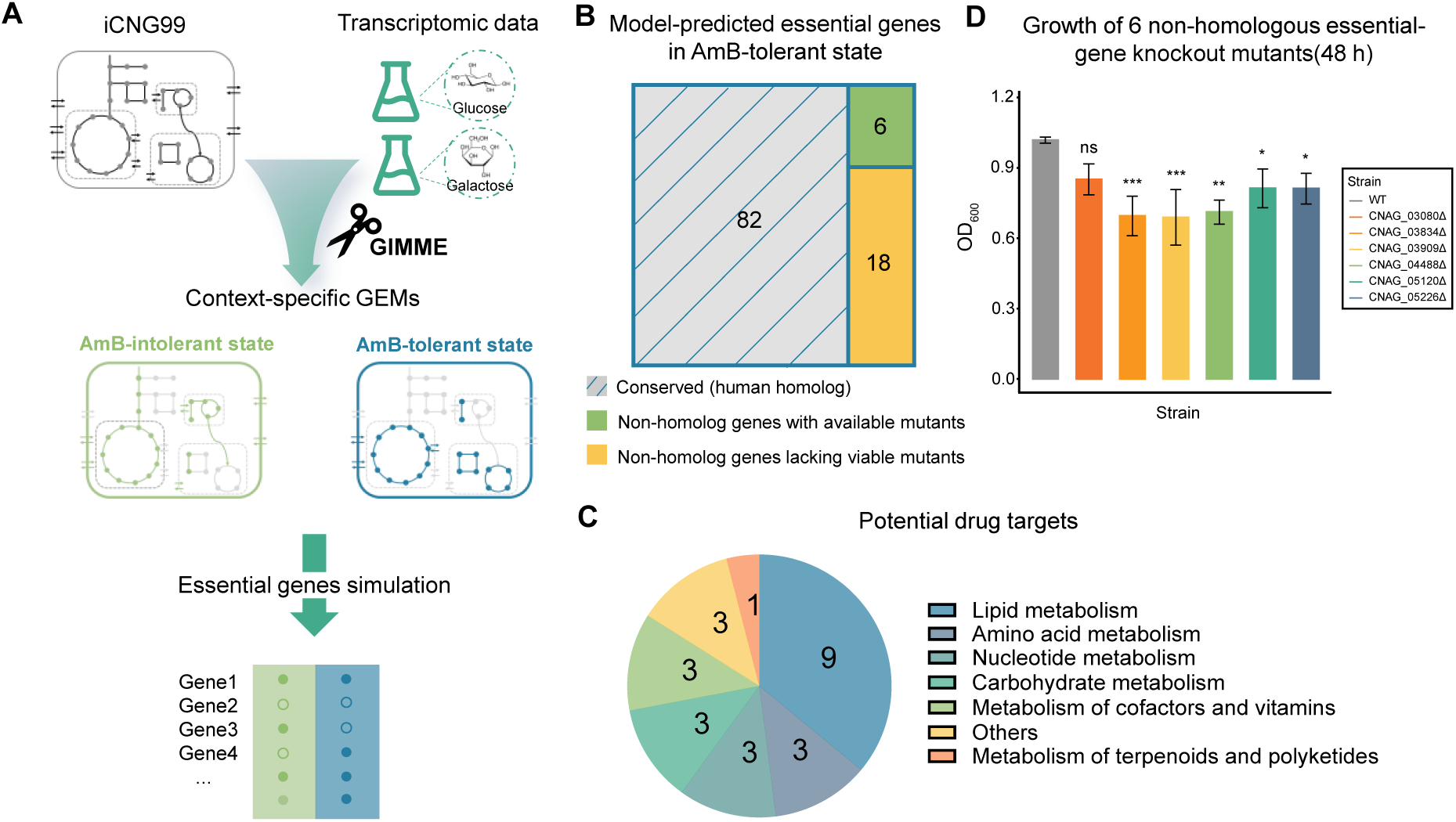
iCNG99 predicts metabolic vulnerabilities in AmB-tolerant state. A. Workflow for predicting essential genes for the AmB-tolerant state. B. Classification of predicted essential genes for the AmB-tolerant state. Predicted essential genes were grouped as conserved (gray), Non-homolog genes with available mutants (green), and Non-homolog genes lacking viable mutants (yellow) based on human homology and mutant library comparison. C. Functional classification of 24 model-predicted essential genes in the AmB-tolerant state that do not have homologs in the human genome. D. OD_600_ at 48 h of six model-only essential-gene mutants under the AmB-tolerant state. Bars represent mean ± s.d. from three biological replicates. Statistical significance at 48 h was determined by Dunnett’s post-hoc test comparing each mutant to the wild-type strain (p.adjust < 0.05, *; p.adjust < 0.01, **; p.adjust < 0.001, ***; p.adjust > 0.05, ns).

## Discussion

In this study, we present iCNG99, a well-curated and experimentally validated GEM of *C. neoformans var. grubii* H99. Previous studies have reconstructed a draft GEM for *C.neoformans* termed iCryptococcus, but systematic benchmarking of its predictive performance remained limited [71]. Successful application of GEMs requires rigorous quality control, comprehensive benchmarking, and experimental validation. Therefore, as iCNG99 has demonstrated high-confidence prediction of essentiality and reproducible characterization of condition-specific metabolic shifts across independent datasets, it extends the previous efforts in modeling *C. neoformans* metabolism and establishes a more robust computational platform for potential translational applications. Metabolic plasticity is central to the ability of *C. neoformans* to survive within the host and under antifungal pressure [1, 72]. Existing metabolic studies on *C. neoformans* have largely focused on individual pathways such as capsule biosynthesis, melanin synthesis, and lipid metabolism [73, 74]. By integrating curated GEM and multi-omics data, iCNG99 was able to provide a systems-level view of how distinct virulence-associated processes involve metabolic shifts between growth and maintenance.

Moreover, the reconstruction strategy of iCNG99 highlights methodological considerations for metabolic modeling of other non-model fungi. *C. neoformans*, as well as other non-model organisms, has incomplete annotation of gene functions and limited characterization of metabolic activities. Therefore, reliance on a single database or computational pipeline can introduce systematic biases that might impair quality of the model. To address these challenges and enhance robustness of the model, we employed an ensemble-based approach throughout the model reconstruction workflow, which integrates multiple resources of pathway annotations, complementary tools for directionality and compartment prediction, together with extensive manual curation and gap-filling. Such strategy could be particularly useful for reconstructing metabolic networks of other pathogenic fungi where high-quality annotations and experimental datasets are missing.

However, several limitations of the iCNG99 need to be acknowledged. First, as a constraint-based metabolic model, iCNG99 focuses on metabolic reactions and does not explicitly include regulatory links or non-metabolic essential functions, which likely contribute to the modest recall observed in prediction of essential genes. Second, many of the GPR associations in the model rely on simplified “OR” logic, which may overestimate the redundancy of metabolic genes and therefore further lower the recall in essentiality prediction. Third, while the model has been extensively optimized through manual curation, annotations for subcellular location of reactions and transport between compartments partly rely on computational predictions that would benefit from experimental validation. Finally, condition-specific flux predictions depend on the assumption that transcript abundance and flux capacity correlate with each other and therefore do not fully capture post-transcriptional regulations and enzyme kinetics. Further refinements of the flux predictions require incorporation of thermodynamic constraints or isotope tracing experiments. Despite these limitations, the high precision of essentiality prediction supports the model’s suitability for predicting metabolic vulnerabilities.

## Data and code availability

All data, code, and models are available at GitHub (https://github.com/sysyboom/iCNG99) and archived on Zenodo (https://doi.org/10.5281/zenodo.18263897). The transcriptomic dataset generated in this study for the *in vivo* versus *in vitro* comparison has been deposited in the Gene Expression Omnibus (GEO) under accession number GSE327336.

## Author contributions

C.F. and Z.D. conceived and designed the study and wrote the manuscript. C.F. performed iCNG99 reconstruction, curation, validation and computational analyses. P.H. and Y.Z. generated and provided experimental data for strain growth and gene knockout assays. W.K. provided the *in vivo* and *in vitro* transcriptomic dataset. X.G. provided metabolomic data under thermotolerant and non-thermotolerant conditions. C.D., B.Z., and L.W. provided guidance and supervision throughout the project.

## Acknowledgements

This research was supported by the National Key Research and Development Program of China (2021YFA0911300 to ZD and BZ), the National Natural Science Foundation of China (12371489 to ZD and 32461160269 to LW), and the Shenzhen Medical Academy of Research and Translation (D2401022 to ZD).

## Tables

**Table 1:**
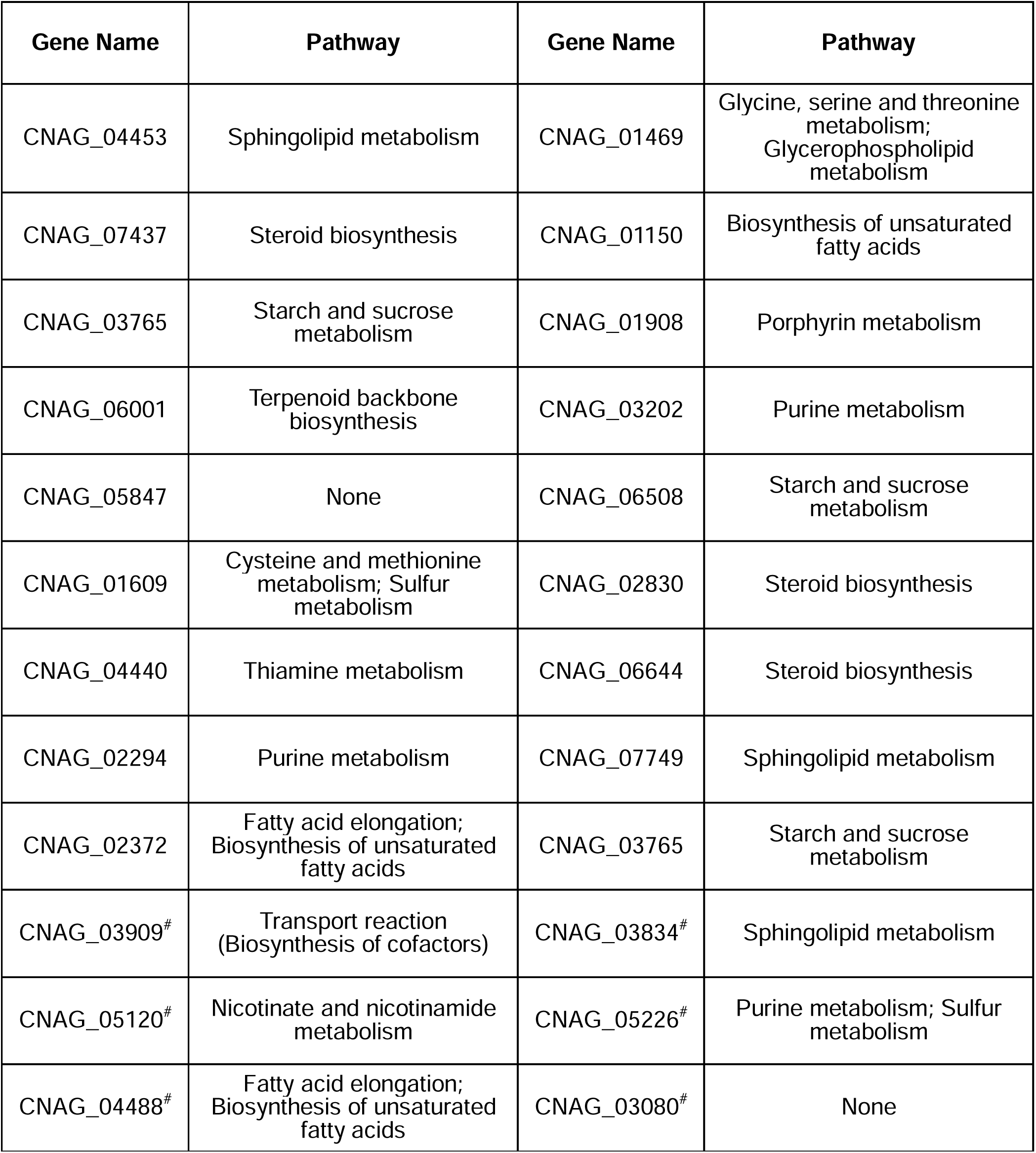
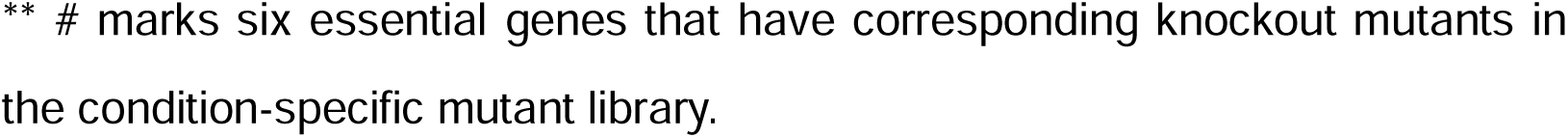
Model-predicted essential genes without human homologs in the AmB-tolerant state.

| Gene Name | Pathway | Gene Name | Pathway |
| --- | --- | --- | --- |
| CNAG_04453 | Sphingolipid metabolism | CNAG_01469 | Glycine, serine and threonine metabolism;<br>Glycerophospholipid metabolism |
| CNAG_07437 | Steroid biosynthesis | CNAG_01150 | Biosynthesis of unsaturated fatty acids |
| CNAG_03765 | Starch and sucrose metabolism | CNAG_01908 | Porphyrin metabolism |
| CNAG_06001 | Terpenoid backbone biosynthesis | CNAG_03202 | Purine metabolism |
| CNAG_05847 | None | CNAG_06508 | Starch and sucrose metabolism |
| CNAG_01609 | Cysteine and methionine metabolism; Sulfur metabolism | CNAG_02830 | Steroid biosynthesis |
| CNAG_04440 | Thiamine metabolism | CNAG_06644 | Steroid biosynthesis |
| CNAG_02294 | Purine metabolism | CNAG_07749 | Sphingolipid metabolism |
| CNAG_02372 | Fatty acid elongation; Biosynthesis of unsaturated fatty acids | CNAG_03765 | Starch and sucrose metabolism |
| CNAG_03909<sup>#</sup> | Transport reaction (Biosynthesis of cofactors) | CNAG_03834<sup>#</sup> | Sphingolipid metabolism |
| CNAG_05120<sup>#</sup> | Nicotinate and nicotinamide metabolism | CNAG_05226<sup>#</sup> | Purine metabolism; Sulfur metabolism |
| CNAG_04488<sup>#</sup> | Fatty acid elongation; Biosynthesis of unsaturated fatty acids | CNAG_03080<sup>#</sup> | None |

\*\* # marks six essential genes that have corresponding knockout mutants in the condition-specific mutant library.

