## Supplementary Fig for "iCNG99: a validated genome-scale metabolic model of *Cryptococcus neoformans strain* H99"

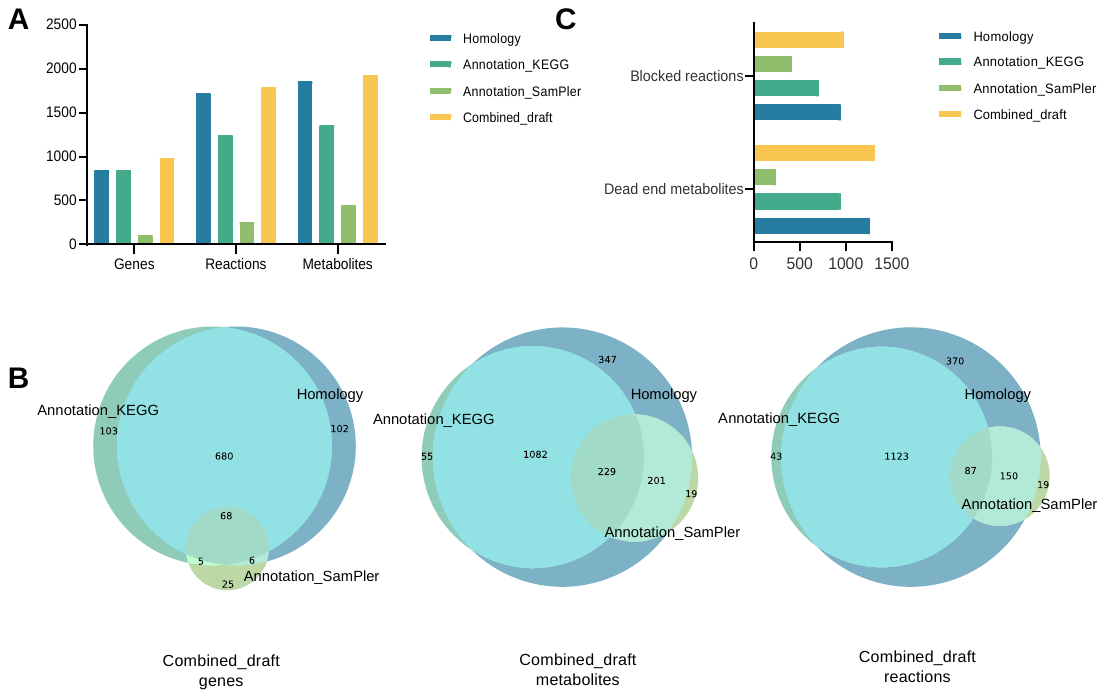

**Supplementary Figure 1: Testing of GEM drafts reconstruction algorithms**

A Comparison of the numbers of genes, reactions, and metabolites in draft metabolic network models generated by different reconstruction methods.

B Comparison of gene, reaction, and metabolite overlaps among draft metabolic network models generated by different methods.

C Statistics of blocked reactions and dead-end metabolites in draft metabolic network models generated by different methods.

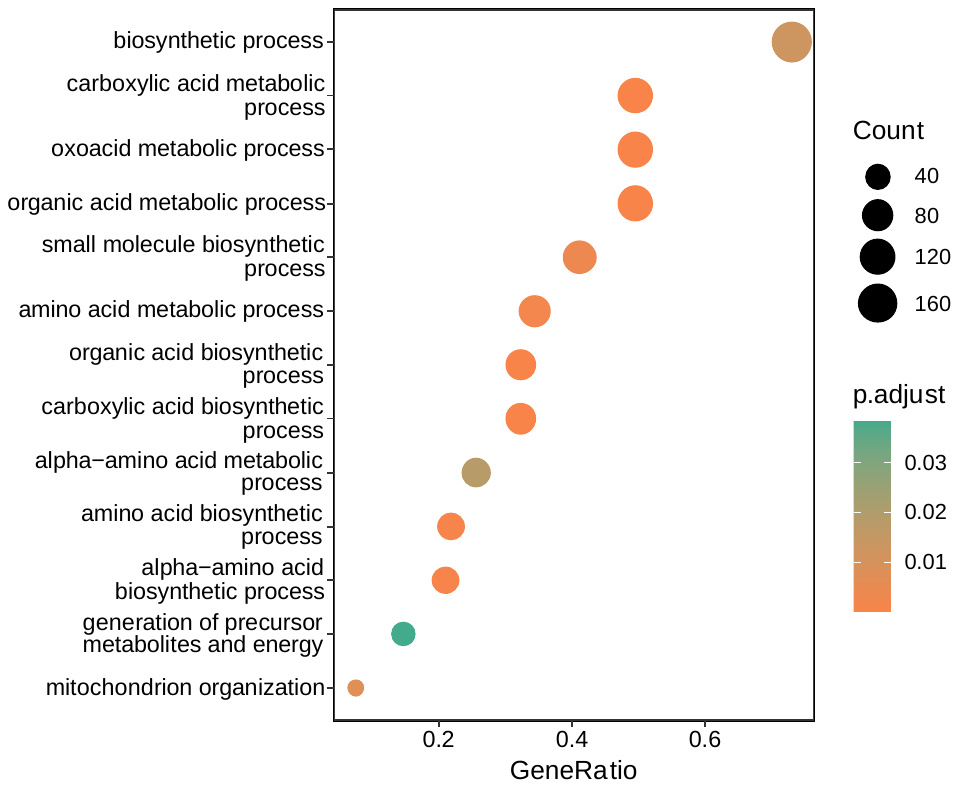

**Supplementary Figure 2: GO enrichment of false-negative (FN) genes**

**Supplementary Figure 3: Correlation analysis of RNA-Seq log_2_ fold changes of *in vivo* relative to *in vitro* conditions derived from two independent transcriptomic datasets.**
